# Absence of the Earth’s magnetic field activates stress-related brain circuits in a mammal

**DOI:** 10.64898/2026.10.04.756554

**Authors:** Li Zhang, Billy Y.K. Lam, Alireza Saeedi, Leoni-Marie Webb, Alessandro A. Felder, Nikoloz Sirmpilatze, Adam L. Tyson, Brittany Hamauei, Runita N. Shirdhankar, Sybille Wolf-Kümmeth, Dori Grijseels, Nasibeh Amini, Sabine Begall, E. Pascal Malkemper

## Abstract

The Earth’s magnetic field is a constantly available cue for animals with a magnetic sense. While magnetic navigation is best studied in migratory birds, a magnetic sense is also found in some mammals. Among these, it is most thoroughly studied in African mole-rats of the genus *Fukomys*, subterranean mammals that navigate in kilometre-long dark tunnel systems. Our understanding of the neural pathways involved in magnetoreception remains incomplete. To address this, we here performed an unbiased whole-brain screen for magnetically induced neuronal activity in the mole-rat brain. We set up MoBI-DisC (Mole-rat Brain Imaging with iDISCO and ClearMap), an experimental pipeline that combines global c-Fos immunolabelling, tissue clearing, light-sheet microscopy, and mapping to a novel 3D brain atlas. To identify brain areas involved in magnetosensation, we exposed mole-rats to a changing magnetic field and compared their brain activity to that of control animals exposed to a near-zero magnetic field. Unexpectedly, we found that the near-zero magnetic field exposure activated many brain areas, predominantly those involved in stress and anxiety responses. These included the amygdala, the bed nucleus of the stria terminalis, the substantia innominata, and the periaqueductal grey. This suggests that magnetic field deprivation acts as a physiological stressor in mole-rats. To test this, we conducted a place-preference test and found that mole-rats avoid entering a zone with a near-zero magnetic field, consistent with the hypothesis that this condition is not neutral for these animals. In summary, we present a novel pipeline for unbiased mapping of neural activity in subterranean mole-rats and uncover evidence of stress and avoidance responses to the absence of an Earth-strength magnetic field.

## Introduction

Many animals from invertebrates to mammals are able to sense the Earth’s magnetic field and use it for orientation (*1*). While the existence of the magnetic sense is behaviourally established beyond doubt, we know little about the peripheral transduction mechanism and central brain circuits that process magnetic cues (*2, 3*). An understanding of the neuronal circuits involved in magnetoreception will enable refinement of mechanistic hypotheses and direct the search specifically towards the elusive primary magnetoreceptive cells (*2*).

Studies in birds using the immediate early genes (IEGs, e.g., c-Fos or ZENK) to map neuronal activity have provided evidence for the involvement of retinal (*4*), trigeminal (*5*), and vestibular circuits (*6, 7*) in magnetic sensing. Among these, the trigeminal pathway has received experimental support in several bird species and also in fish (*5, 8–10*). This has led to the hypothesis that magnetoreceptors based on magnetite are associated with nerve endings of the ophthalmic branch of the trigeminal nerve, but several attempts to find such sensory structures have failed to provide conclusive evidence (*11–13*). The variety of circuits reported may reflect multiple magnetosensory mechanisms, differences in experimental design, or species-specific variation, highlighting the need for further studies across taxa.

Among mammals, magnetic orientation has been demonstrated in bats (*14*) and rodents (*15–17*), but data on magnetosensory brain areas are limited to a single species: the subterranean Ansell’s mole-rat (*Fukomys anselli*) (*18*). These highly social bathyergid rodents inhabit convoluted underground tunnel systems that are constantly remodelled and extended (*19*). They possess a magnetic polarity compass that works in total darkness (*20*) and is unaffected by weak radiofrequency magnetic fields (*21*), consistent with single-domain magnetite receptors as proposed for the trigeminal system of birds. The absence of directional preferences after enucleation or local anaesthesia of the cornea provided support for a localisation of these receptors in the eye (*22, 23*), but an extensive multimodal imaging screen could not identify magnetite (*13*). Hypothesis-driven c-Fos mapping has revealed magnetic responses in the superior colliculus (SC) and a number of brain regions of the rodent navigation circuit, including the hippocampus (HC), the entorhinal cortex (EC), and the anterodorsal thalamic nucleus (ADN) (*24, 25*), supporting a role of magnetic cues in spatial navigation. However, it is unclear how these higher-order brain regions receive magnetic information.

We here performed an unbiased, global screen for magnetically induced neuronal activity in the Ansell’s mole-rat brain. Recent advances in tissue clearing and volumetric imaging of fluorescently labelled immediate early genes (IEGs) now enable cellular-resolution mapping of sensory-evoked neuronal activity throughout the brain, providing an unbiased view of where neural responses occur (*6, 26*). We used such an approach to compare neuronal activity in mole-rats exposed to magnetic stimuli previously shown to engage the magnetosensory system in birds (*5, 9*). Unexpectedly, this revealed that exposure to a near-zero hypomagnetic field recruited activity across more brain regions than slow changes in magnetic field strength and direction, with responses predominantly observed in stress-related nuclei.

## Results

### MoBI-DisC: A pipeline for whole-brain IEG mapping in mole-rats

To map IEG expression across the mole-rat brain in an unbiased manner, we adapted and extended the existing ClearMap 2.0 software for mice (*26, 27*) and used it to develop a package which we named MoBI-DisC (**Mo**le-rat **B**rain **I**maging with **DIS**CO and **C**learMap). The procedure integrates whole-brain immunohistochemistry, solvent-based iDISCO+ tissue clearing (*28*), light-sheet microscopy, machine learning-based 3D cell detection (*26*), and registration to a newly generated mole-rat 3D brain atlas for voxel-wise statistical analysis, which is available through the BrainGlobe Atlas API (*29*)(see Methods; **Fig. 1**). MoBI-DisC enables the automated identification of brain areas that differ in IEG expression between two or more treatment groups. It includes a permutation-based correction for multiple testing and performs back-projection of significant clusters from the cluster-corrected *p*-value map into the original 3D image space. This allows extraction of raw counts and provides direct visual confirmation that the statistically significant voxels corresponded to genuine, anatomically coherent groups of c-Fos-positive cells, thereby mitigating the risk of false positives arising from registration artefacts or sparse cell distributions.

**Figure 1.**
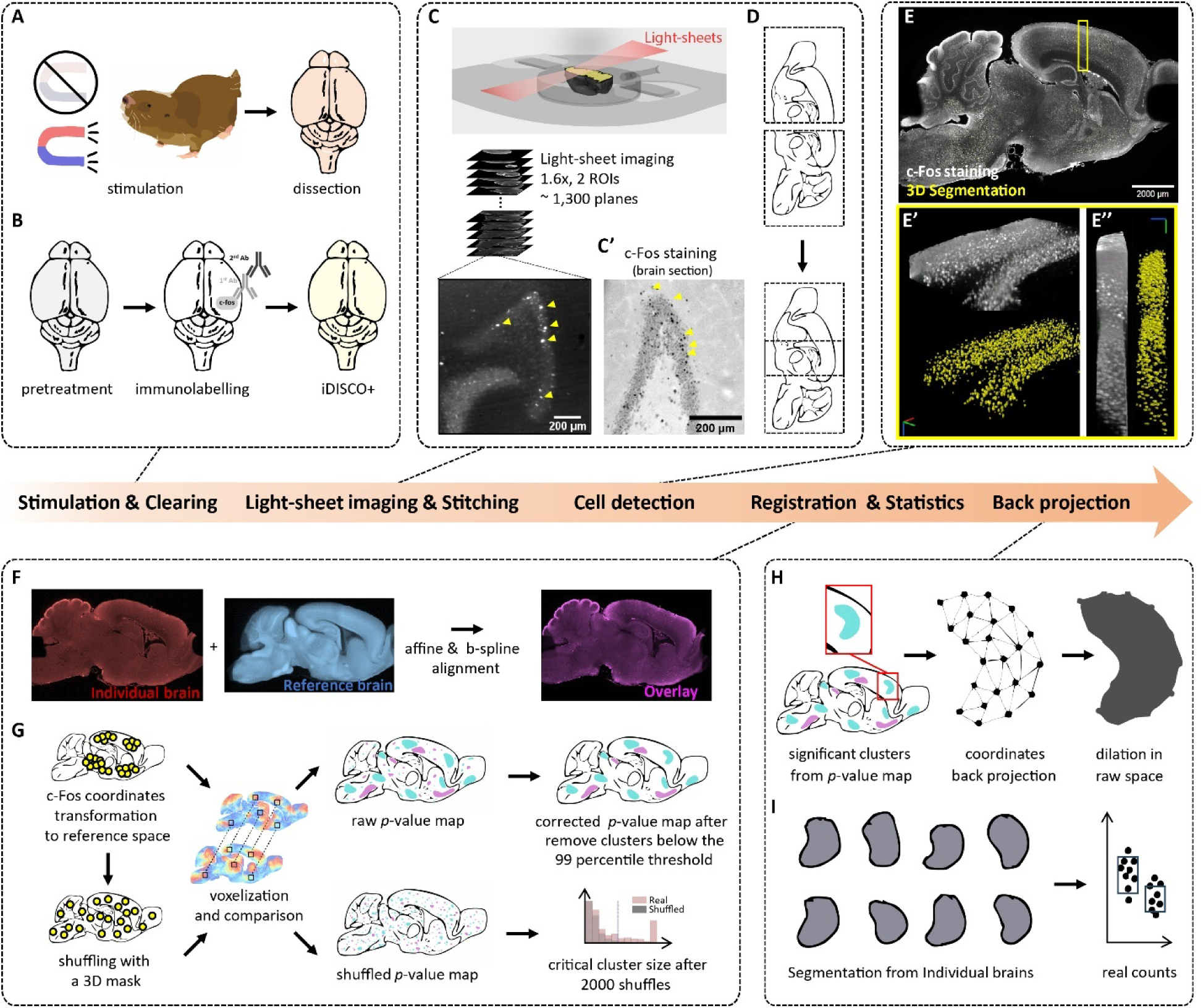
| The MoBI-DisC pipeline for unbiased whole-brain neural activity mapping in mole-rats. (**A**) Sensory stimulation followed by brain collection after transcardial perfusion. (**B**) Whole-brain immunohistochemistry followed by solvent-based tissue clearing. (**C**) Light-sheet imaging of cleared brains. Images were acquired in the sagittal plane, starting at the cortical surface. Yellow arrowheads in a virtual section indicate c-Fos-positive nuclei in the hippocampal dentate gyrus in comparison to (C’) traditional DAB staining of c-Fos on a brain section. (**D**) Imaging strategy. Two fields of view were imaged and digitally stitched to cover the rostro-caudal axis. (**E**) Machine learning–based 3D cell detection using Ilastik. Example plane showing overlay of c-Fos staining (grey) and Ilastik segmentation (yellow) (E), 3D rendering of a cortical subvolume (E’), and its sagittal view (E’’). Orientation cross: red (rostral-caudal), green (dorsal-ventral), blue (medial-lateral) in (E’) and (E’’). (**F**) Image registration and (**G**) *p*-value map generation with customised ClearMap (*26*). Individual brains were registered to a mole-rat 3D reference atlas via Elastix (F), followed by voxelization, statistical analysis, and a permutation-based multiple comparison correction of the *p*-value map (G). (**H, I**) Back-projection of significant clusters from the *p*-value map into the 3D raw data space for post hoc validation and extraction of raw counts. Coordinates of a representative cluster were back-projected and dilated in raw space (H), with individual dilated datasets shown for comparison (I).

### An anatomical 3D mole-rat reference brain atlas

To enable differential analyses across brains, we constructed a three-dimensional average mole-rat brain template with a resolution of 20 μm x 20 μm x 20 μm from a cohort of 43 light-sheet scanned brain hemispheres (**Fig. 2A; Supplementary Fig. 3**). We manually annotated 33 major brain areas in this template that we identified based on gross anatomical features, aided by histological 2D atlases of the mole-rat brain (**Fig. 2B, C; Supplementary Fig. 4**) (*30, 31*). To assess the accuracy of the atlas mapping, we performed an anterograde neural tracing experiment in which we injected fluorescently labelled Cholera toxin subunit B (CTB) into the mole-rat eye, which labels known retinal projections to primary subcortical visual nuclei in the brain (**Fig. 2D, E**) (*32*). Registration of the cleared, light-sheet imaged brains to our atlas revealed a strong overlap between the CTB-labelled axon terminals and annotated subcortical visual targets, including the suprachiasmatic nucleus (SCN), lateral geniculate nucleus (LGN), olivary pretectal nucleus (OPN), and superficial layer of superior colliculus (SCs) (**Fig. 2F–J**). Together, these findings confirm the accuracy of the 3D mole-rat brain atlas for anatomical mapping, at least for the relatively small (and therefore challenging) nuclei we examined here.

**Figure 2.**
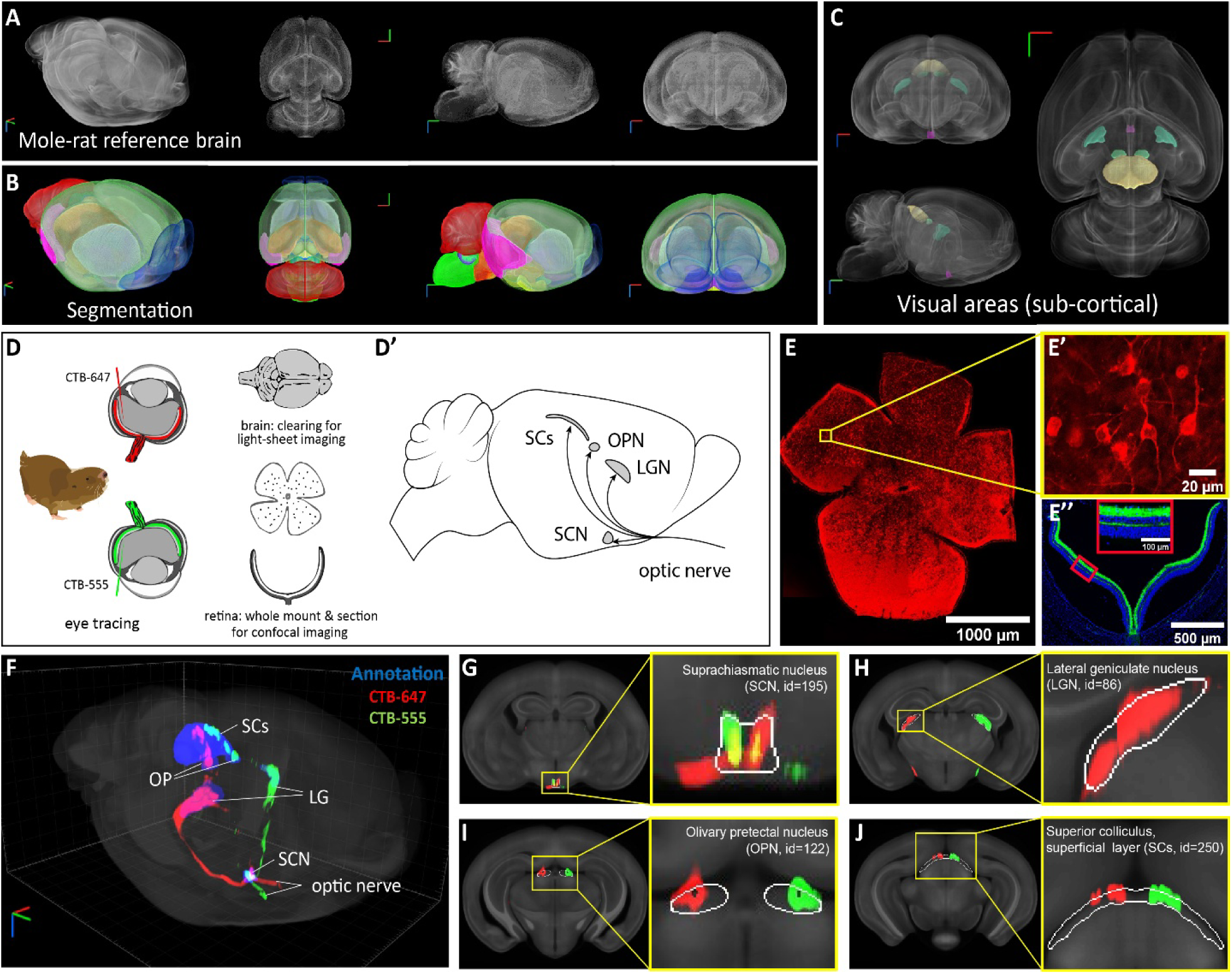
| A 3D mole-rat brain atlas allows to map target brain regions from visual tracing. (**A**) 3D projections of the reference brain viewed from multiple angles (from left: rostral lateral oblique view, horizontal, sagittal, and coronal). (**B**) Corresponding projections of the annotated atlas. (**C**) Visualisation of subcortical visual structures within the atlas framework. (**D**) Overview of the tracing experiment. CTB was bilaterally injected into the eye. We analysed the retinas as flat mounts or sections, and the brain with whole-brain clearing and MoBI-DisC. (D′) Schematic of the visual pathway showing the expected target brain regions: suprachiasmatic nucleus (SCN), lateral geniculate nucleus (LGN), olivary pretectal nucleus (OPN), and superficial layer of superior colliculus (SCs). (**E**) Representative fluorescence images of CTB-positive cells in whole mounts (E, E′) and sections (E″). (**F**) Tracing results registered to the annotated reference atlas. (**G–J**) Overlays of retinal projections with annotated subcortical targets (white boundaries). Orientation cross in (A–C) and (F): red (medial–lateral), green (rostral–caudal), blue (dorsal– ventral).

### Magnetically induced c-Fos expression in the mole-rat brain

We next used MoBI-DisC to investigate brain-wide neural responses to magnetic field changes. Mole-rats were habituated 30 min daily to sit in a small bucket in total darkness. On the fourth day, the animals were exposed to either a changing magnetic field (CMF) or a near-zero hypomagnetic field (HMF, <1 µT, ∼ 2% of the Earth’s magnetic field intensity in Germany) for 72 minutes before brain collection and processing through the MoBI-DisC pipeline (**Fig. 3A, B**). The total number of c-Fos-positive cells per hemisphere was not significantly different between the CMF (median = 144,022 cells) and HMF (median = 181,327 cells) groups (Mann-Whitney test, U = 32, n = 10 per group, *p* = 0.1903; **Fig. 3C**), indicating that the magnetic stimulation did not induce global changes in neural activity. Furthermore, manual ethograms revealed no differences in behaviour of the animals in both treatment conditions (**Supplementary Fig. 4).**

**Figure 3.**
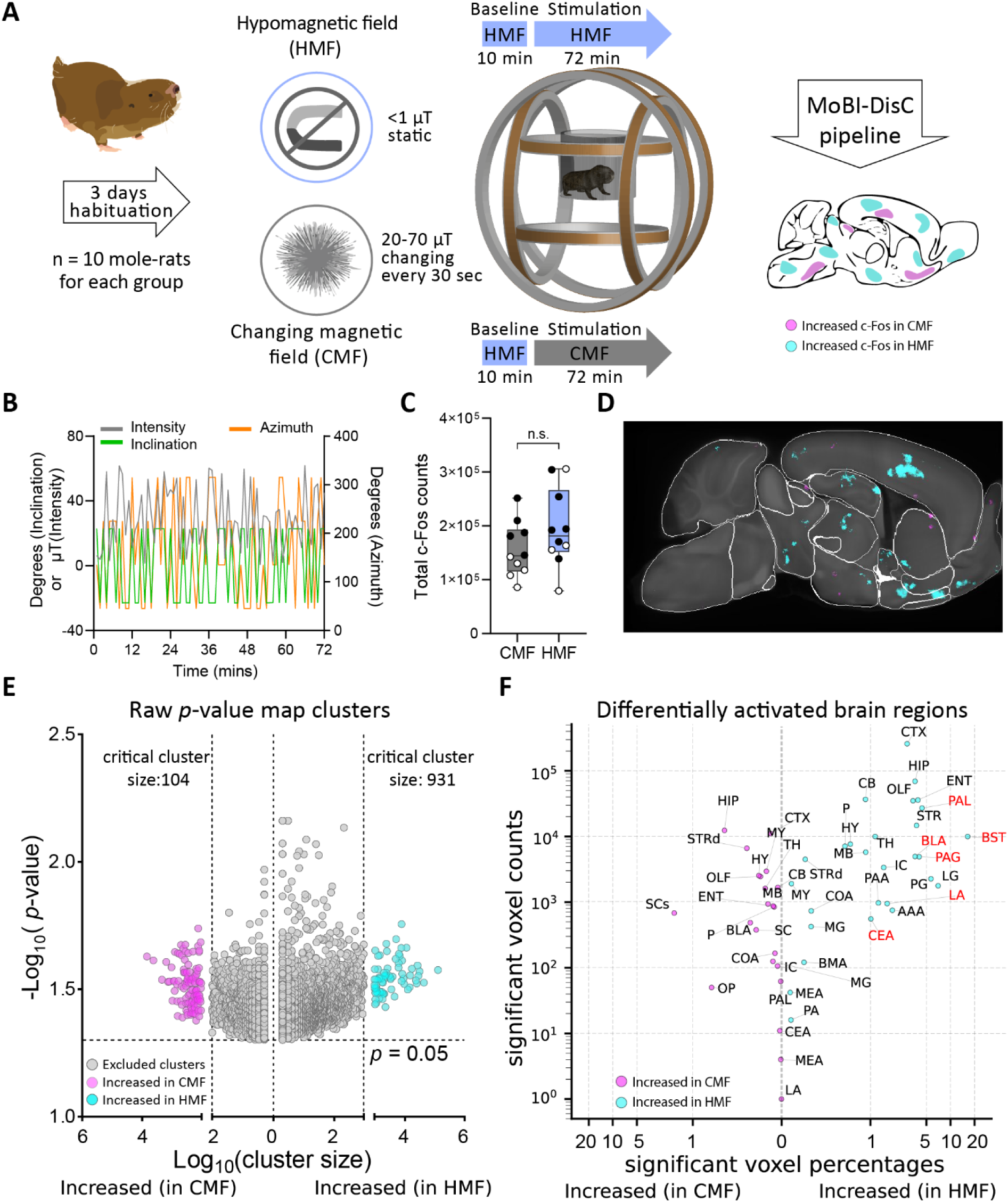
| Global mapping of mole-rat brain areas activated by magnetic stimulation. **(A)** Experimental design of the magnetic-field stimulation paradigm, followed by whole-brain c-Fos mapping with the MoBI-DisC pipeline. (**B**) Representative magnetic field intensity traces along the x-, y-, and z-axes of a magnetometer, illustrating the temporally fluctuating field in the CMF condition. (**C**) Total c-Fos positive cell counts in brain hemispheres of the CMF and HMF groups (Mann-Whitney test, U = 32, n=10 per group, *p* = 0.1903, n.s. = not significant). Animals from two independent experimental cohorts are shown as open and filled dots. (**D**) Example sagittal plane of the voxel-wise *p*-value map with many voxels with increased c-Fos counts in the HMF group (turquoise). (**E**) Permutation-based correction for multiple testing of significantly different clusters. The plot shows the size distribution of significant clusters from the raw *p*-value maps (threshold *p* < 0.05). Each dot represents a spatially contiguous voxel cluster; vertical dashed lines denote empirically derived critical cluster sizes corresponding to the cluster-based correction thresholds (see also **Supplementary** Fig. 5). (**F**) Total and relative differences in differentially activated voxels of all brain regions annotated in our mole-rat brain atlas. The brain regions highlighted in red belong to circuits associated with stress, fear, and anxiety. An overview of the brain region abbreviations can be found in **Supplementary** Fig. 3).

Voxel-wise analysis followed by cluster-based multiple comparison correction (**Supplementary Fig. 5**) revealed a distinct pattern of localised activation. Surprisingly, only a few brain regions showed a significant fraction of voxels in which c-Fos expression was higher in the CMF than in the HMF group (**Fig. 3D–F, Supplementary Table 1**). Most notably, we found a localised increase in the dorsal subiculum within the hippocampal formation (n = 10 per group, U = 11, p_adj_= 0.003; Supplementary Fig. 6), a region known as a hub for navigation-related information (*33*).

Conversely, we identified numerous discrete clusters across the brain that exhibited higher c-Fos density in the HMF group (**Fig. 3D–F, Supplementary Table 1**). These were concentrated on subcortical brain regions canonically associated with stress, fear, and anxiety (**Fig. 4A**), such as the posterior basolateral amygdala (BLAp; Mann-Whitney tests with FDR correction (Q=5%), n = 10 per group, U = 22, p_adj_ = 0.012), the central amygdala (CeA; n = 10 per group, U = 14, p_adj_ = 0.003), lateral amygdala (LA; n = 10 per group, U = 24, p_adj_ = 0.015), and the bed nucleus of the stria terminalis (BST; n = 10 per group, U = 12, p_adj_ = 0.003). Elevated activity was further observed in downstream regions of the amygdalar circuitry, including the periaqueductal grey (PAG; n = 9 for CMF, n = 10 for HMF, U = 24.5, p_adj_ = 0.026), substantia innominata (SI; n = 10 per group, U = 17, p_adj_ = 0.006), and the nucleus accumbens (ACB; n = 10 for CMF, n = 9 for HMF, U = 17, p_adj_= 0.009) (**Fig. 4B–H**). Taken together, these findings show that the HMF, intended as a baseline control stimulus, activated core brain circuits that coordinate stress and defensive responses in the absence of obvious behavioural changes, indicating that HMF exposure exerted a mild aversive effect.

**Figure 4.**
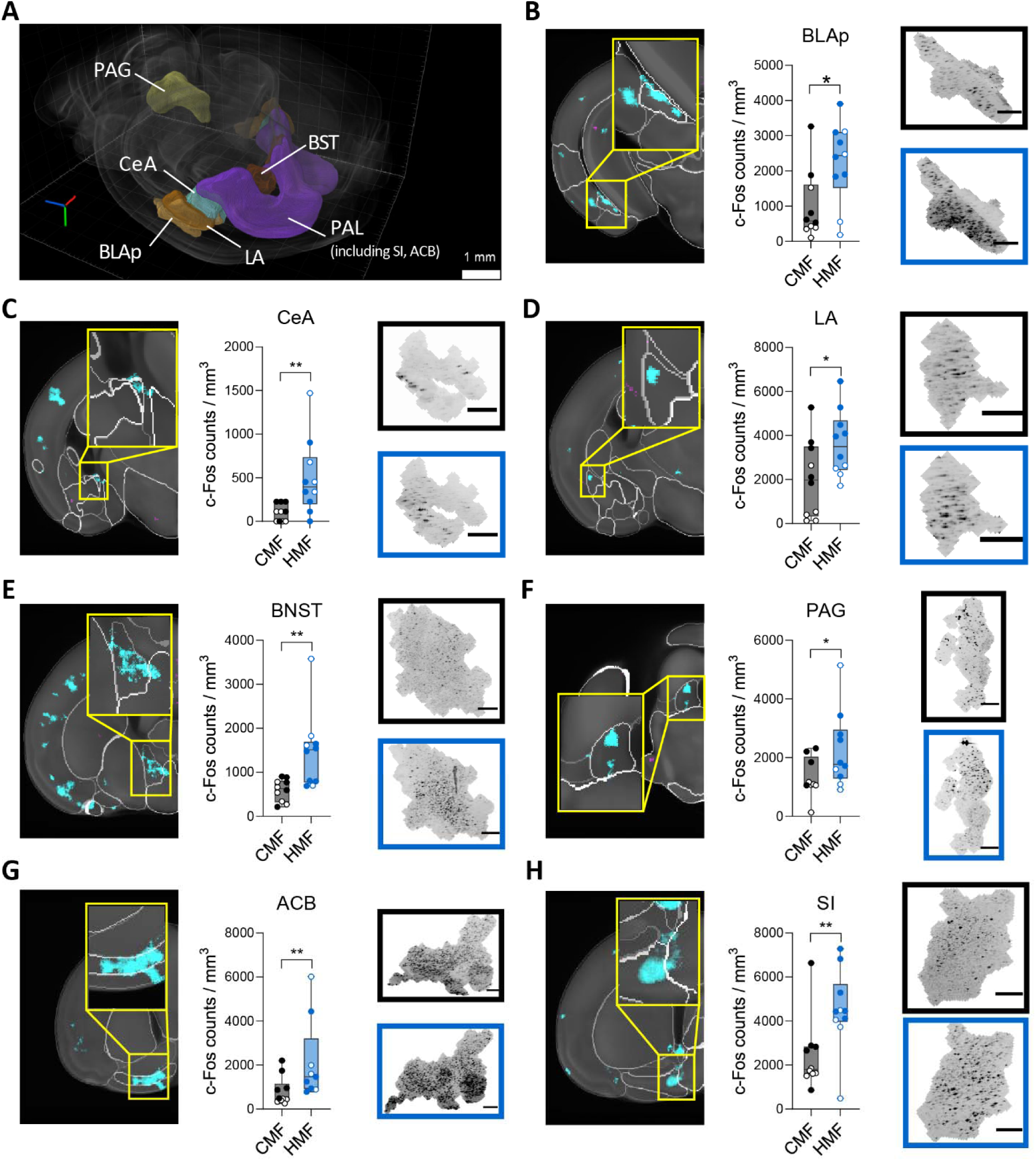
| Stress-associated brain regions with increased c-Fos expression after exposure to HMF. (**A**) 3D visualisation of brain regions showing significant differences between CMF and HMF stimulation, mapped onto the mole-rat reference atlas. Orientation cross: red (rostral–caudal), green (dorsal–ventral), blue (medial–lateral). (**B–H**) Increased c-Fos–positive cell density under HMF in seven representative brain regions. Left, single planes of the voxel-wise *p*-value maps with insets; middle, box plots comparing cell densities; right, back-projection examples for each region, the examples shown correspond to the dataset entry closest to the mean value of each group from the same experiment. Statistical significance was assessed using Mann–Whitney tests with FDR correction (Q = 5%), * *p_adj_*< 0.05, ** *p_adj_*< 0.01 and exact *p_adj_* values are reported in the main text. n = 10 mole-rats per group; solid and open data points represent two independent repeats. Scale bars: 200 μm. BLAp: Basolateral Amygdala, posterior part; CeA: Central Nucleus of the Amygdala; LA: Lateral Amygdala; BNST: Bed Nucleus of the Stria Terminalis; PAG: Periaqueductal Gray; PAL: Pallidum; SI: Substantia Innominata; ACB: Accumbens Nucleus.

### Mole-rats avoid a hypomagnetic field anomaly in a place preference test

Based on the neural activation patterns indicative of a stress-like state induced by HMF exposure, we hypothesised that mole-rats perceive the lack of a magnetic field as aversive and consequently exhibit active avoidance behaviour to HMF. To test this hypothesis, we designed a closed-loop behavioural paradigm in which the instantaneous position of the animal within the arena dynamically controlled the local magnetic field intensity, thereby generating spatially defined “virtual” zones of HMF (near zero-field anomalies, **Fig. 5A-D**). The background magnetic field intensity was 44.71 µT (SD 0.02, range 44.68–44.76 µT; n = 473 measurements) at the arena centre and 1.45 µT (SD 0.73, range 1.08–4.88 µT; n = 713 measurements) at the virtual magnetic anomaly (HMF) (**Fig. 5D**). Using this paradigm, we performed a simple place preference test in a linear track with two non-magnetic novel objects (e.g., laboratory plastic 96-well plates) positioned at opposite ends of the arena (**Fig. 5B**). The occupancy within predefined regions of interest (ROIs) centred on each object, defined as the cumulative time spent in proximity to either object over the course of 20 min trials, was taken to assess the preference of the animals (**Fig. 5E**).

**Figure 5.**
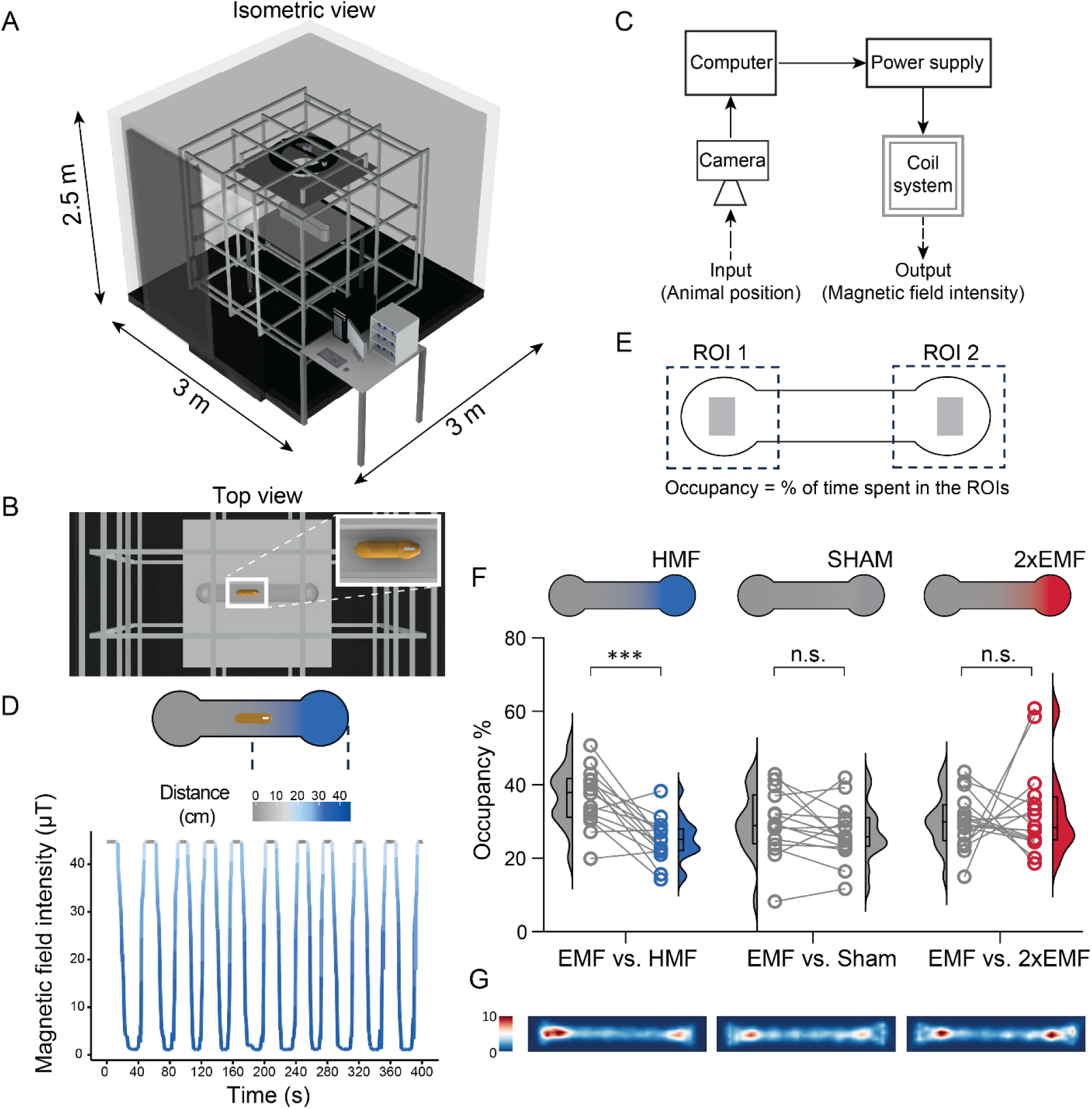
| Mole-rats avoid a hypomagnetic field anomaly in a place preference test. (**A**) Experimental setup. A linear track was placed on a table in the centre of a 3D Merritt-coil system. The experiments were performed in an isolated room that shielded sound, light, vibrations, and oscillating magnetic fields. (**B**) The animal was allowed to inspect two non-magnetic objects placed at the end of the track for 20 min. (**C**) Scheme of the closed-loop magnetic field control, which created a local magnetic field anomaly by varying the magnetic coil system output based on the animal’s position. (**D**) Representative magnetic field measurements as a function of position over time. (**E**) Relative occupancy was quantified based on the time spent in ROI centred on the objects. (**F**) Paired comparisons of occupancy within two ROIs across three experimental conditions: EMF vs HMF (left), EMF vs Sham (middle), and EMF vs 2xEMF (right). Magnetic field intensities are colour-coded. n = 15/16 mole-rats per experiment. Statistical significance was assessed using paired two-tailed t-tests for EMF vs HMF, t (14) = 4.62, p = 0.0004; EMF vs Sham, t (14) = 1.23, p = 0.240; EMF vs 2xEMF, *t* (15) = 0.518, *p* = 0.612. Individual data points are shown alongside a violin plot to show the distribution of each data group. (**G**) Averaged occupancy maps of the three conditions (arbitrary units).

Mole-rats spent significantly less time in the HMF ROI than in the EMF (paired t-test, *t*(14) = 4.67, *p* = 0.0004; Fig. 5C, left), demonstrating a clear preference to avoid the HMF (**Fig. 5D, left**). Importantly, this avoidance was specific to the hypomagnetic condition, as control experiments showed no significant preference between EMF and Sham (with current flow but no magnetic field change, details see Methods; paired t-test, *t* (14) = 0.123, *p* = 0.240; **Fig. 5F, middle**) or between EMF and a magnetic field with doubled intensity (2xEMF; paired t-test, *t* (15) = 0.518, *p* = 0.612; **Fig. 5C**, **Fig. 5F, right**). The distribution of individual animal performances and avoidance indices further corroborated consistent, specific avoidance of HMF across the cohort. The avoidance index for the HMF condition was significantly greater than zero (one-sample t-test, *t* (14) = 4.62, *p* = 0.0004), whereas the indices for the Sham (*t* (14) = 1.04, *p* = 0.316) and 2xEMF (*t* (15) = 0.287, *p* = 0.778) conditions were not (**Supplementary Fig. 7**). In summary, these behavioural experiments show that mole-rats actively and specifically avoid near-zero magnetic field anomalies, which is in line with the finding of activated stress-related brain areas after forced exposure to HMF.

## Discussion

We aimed to identify neuronal circuits involved in magnetoreception and performed an unbiased whole-brain screen for neuronal activation induced by magnetic field changes in the mole-rat brain. To do so, we mapped global c-Fos expression patterns after exposure to a CMF and compared these to a control group exposed to an HMF, using a novel open-source 3D reference brain atlas. Unexpectedly, the strongest and most widespread neuronal response was observed not after exposure to the CMF, but after exposure to the nominally neutral HMF. This finding fundamentally changes how the HMF condition should be interpreted: rather than representing the absence of a magnetic stimulus, it appears to constitute a biologically salient sensory perturbation. The neuronal activation patterns are consistent with the mole-rats detecting the absence of the Earth’s magnetic field, presumably via their magnetoreceptors. Behavioural data support the hypothesis that mole-rat magnetoreceptors are based on single-domain magnetite particles coupled to a mechanosensory cell (*13, 20*). Given that the Earth’s magnetic field is a weak stimulus, it is assumed that the magnetoreceptive organ contains an array of sensors to improve the signal-to-noise ratio through spatio-temporal averaging (*34*). One possible explanation is that HMF produces stochastic activation across the proposed magnetite-based receptor array. If individual receptors are differently oriented in the absence of an external field, random subsets of receptors could be activated at any given time, producing aggregate input to downstream nuclei comparable to that elicited by the CMF. Such shared activation would be invisible to our differential c-Fos analysis.

The neuronal activation patterns support a model in which mole-rats sense the absence of the Earth’s magnetic field (**Fig. 6**). The fact that many of the strongly activated regions belonged to circuitry involved in stress-related responses further supports a negative valence for this sensation. The activation of SI and PAG in our mole-rats further indicates that the animals were aroused (*35*). We did not observe differences in behaviour between the treatment groups, however, pointing towards a mild form of anxiety or novelty response, known to activate similar circuits in other rodents. We also did not observe increased head-scanning, a behavioural response to HMF that has been reported in migratory birds (*36*). However, video quality limited the behavioural detail we could assess, and our basic manual ethogram may have missed other, more subtle behavioural responses to HMF.

**Figure 6.**
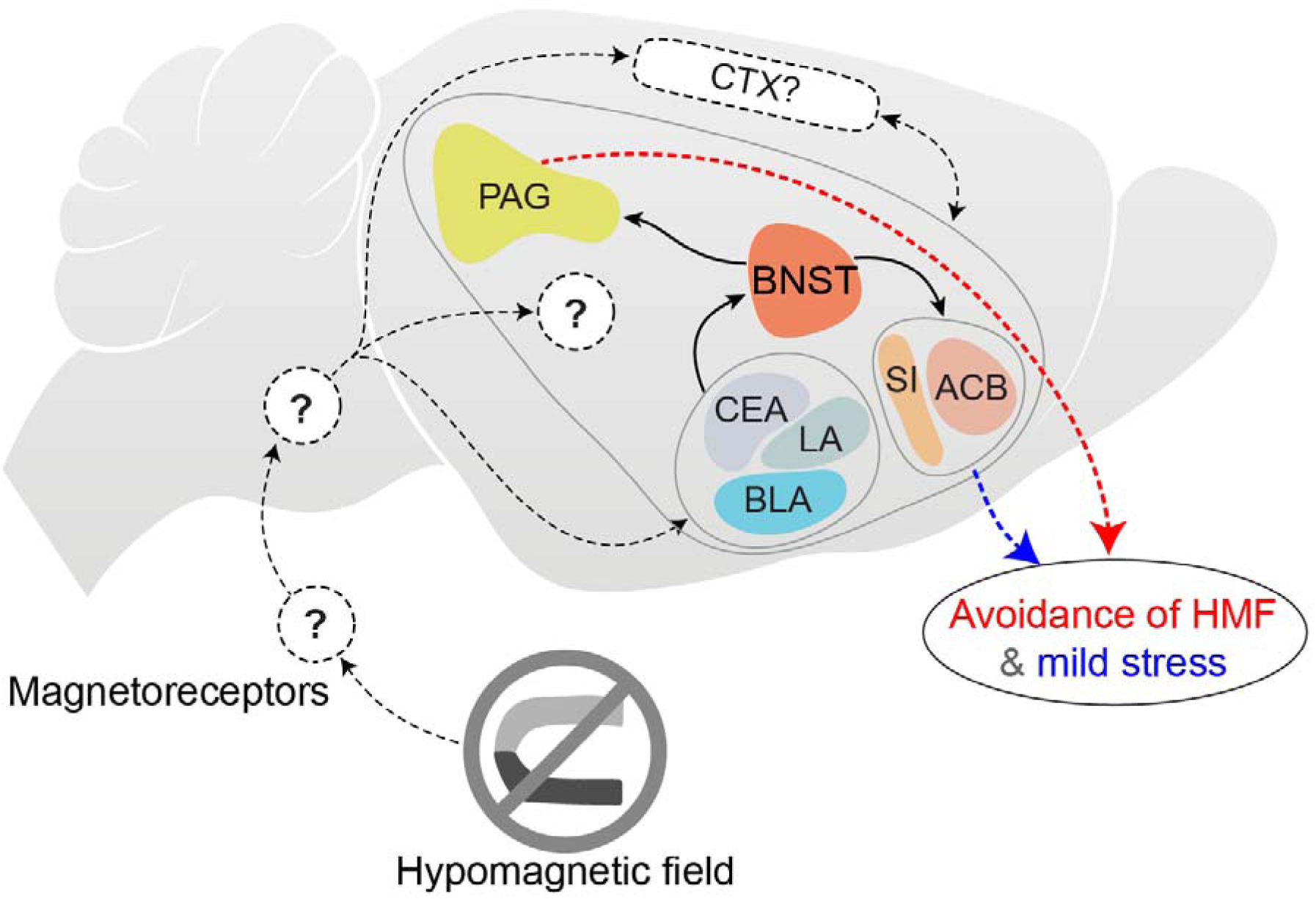
| Model of the involved circuitry in the response to a near-zero magnetic field in mole-rats. The HMF either activates intensity-encoding magnetoreceptors or randomly activates directionally sensitive compass cells. This information activates the central stress-, arousal-, and anxiety-related brain areas via a series of unknown brain areas. This leads to mild stress and corresponding avoidance behaviour. ACB: nucleus accumbens, BLA: basolateral amygdala, BNST: bed nucleus of the stria terminalis, CEA: central amygdala, CTX: cortex, LA: lateral amygdala, PAG: periaqueductal grey, SI: substantia innominata

In rats, exposure to novel environments increases c-FOS expression in several amygdalar nuclei, including the basolateral, medial, and lateral subdivisions (*37–39*), as well as the BNST (*39*). Environmental novelty and stress are also known to activate neurons in motivational areas, including the ACB (*40, 41*). In our experiment, however, the mole-rats had been habituated to the environment, and therefore, the response was unlikely due to novelty alone. Rather than the environment itself being novel, the sudden loss of a normally persistent sensory cue may have rendered the familiar environment perceptually novel. In mice, sensory changes as little as a novel taste added to the drinking water led to increased IEG expression in the amygdala (*42*). While our mole-rats have experienced changes in the local magnetic field in their home cages and during transport to experimental setups (e.g., door frames are inherently magnetic and the metals used in the construction of the building induce anomalies in the range of tens of µT in the holding facilities), they most likely never experienced a near-zero magnetic field in their entire life. The active avoidance of near-zero-field areas in the place preference experiments further supports this hypothesis. While mole-rats are notoriously curious and readily inspect objects, they did so less when the object was placed inside an HMF anomaly. Increasing the magnetic field around the object, in contrast, did not lead to an avoidance behaviour. A study with several mole-rat species and laboratory mice reported that these animals inspected magnetic objects longer than non-magnetic controls (*43*). Except for two animals, we did not observe this behaviour here. On average, the increased field did not make the object more attractive, a difference which could be explained by the intensity of the magnetic anomalies used in both studies. Here, we only doubled the Earth’s magnetic field intensity to ∼90 µT, whereas Malewski et al. (43) used strong magnets with field strengths in the mT range, which are unlikely to be encountered in animal facilities and are therefore novel to the animals. Taken together, these findings thus suggest that mole-rats perceive the absence of an Earth-strength magnetic field as a novel, unexpected sensory deprivation.

We show here, to our knowledge, for the first time, that a mammal shows an active avoidance reaction to the absence of a magnetic field. However, responses to HMF have been reported in many biological systems, from single cells to mammals (*44*). Our findings of behavioural reactions to HMF might be particularly relevant with respect to findings in other rodents. Most comparable to the findings reported here, an increase in anxiety-related behaviours, as assessed in open field and elevated plus-maze tests, was reported for male C57 mice after 72 h of HMF exposure (*45*). Long-term HMF exposure also affects hippocampal neurogenesis and learning in laboratory mice, and researchers linked this effect to reactive oxygen species levels (*46, 47*). Another study found that after exposing laboratory mice to a null field for 1 h a day in a mu-metal chamber, a significant reduction of nociception was detected in the hot-plate test (*48*), an effect that was abolished by the reintroduction of weak magnetic fields (*49*). To date, there is no mechanistic explanation for this analgesic effect. We observed activation of the PAG in mole-rats after HMF exposure, a brain region centrally involved in pain modulation and endogenous opioid signalling. The activation of the PAG therefore raises the possibility that magnetic-field deprivation can influence nociceptive processing. Although this link remains speculative, it provides a testable hypothesis for revisiting the analgesic effects of HMF in mice, particularly in light of evidence for magnetic orientation in this species (*17, 50, 51*).

What did we learn about the magnetic sense of mole-rats from these experiments? First, we can conclude that the magnetic field appears to be a salient environmental cue even in the absence of a navigational task. Second, we learn that, similar to other sensory modalities, magnetic information appears to be distributed across many higher brain areas. This is in line with an earlier c-Fos study in the same species, which found significant magnetic field responses across many brain regions of the rodent navigation circuit (*24*). Third, the animals reacted to the magnetic field changes under IR illumination, which further supports a light-independent mechanism of magnetoreception that differs from the magnetic compass in migratory birds (*21*).

Finally, we also detected some brain areas in which IEG expression was higher after CMF exposure. These included parts of the subiculum and the superior colliculus. Based on our experiments, we cannot tell whether these areas responded with increased activity to CMF or were inhibited by HMF. Encouragingly, however, both of the brain areas have been previously reported to react to magnetic field changes in the same species of mole-rats (*24, 25*). These brain regions therefore represent promising targets for electrophysiological investigations (*52*). The upstream circuits through which the magnetic information reaches these centres should be the focus of future studies. The MoBI-DisC pipeline for whole-brain activity mapping introduced here will facilitate such studies. An important methodological implication of our findings is that HMF cannot be considered a neutral control condition in studies of magnetoreception. Future experiments should therefore compare magnetic stimulation against the Earth’s static field rather than against near-zero-field conditions. To control for self-initiated magnetic stimulation by moving in the static field, such experiments should be performed with head fixation or anaesthesia to maximise control over the magnetic stimulus.

In summary, we present evidence that mole-rats perceive the absence of the Earth’s magnetic field. We show that the removal of this normally constant environmental stimulus, the Earth’s magnetic field, activates stress-related brain circuits and triggers active avoidance behaviour. Our findings demonstrate that the Earth’s magnetic field constitutes a biologically significant sensory stimulus in mammals.

## Materials and methods

### Animals

Adult mole-rats (*Fukomys anselli*) of both sexes were used in this study (see Supplementary Table 2). These animals were maintained under a 12 h light/dark cycle at 20–22 °C with ad libitum access to solid food (potatoes, carrots, and grains). The mole-rats were randomly assigned to experimental groups; experimenters performing outcome measurements were blinded to group allocation where feasible. All animal experiments were performed in accordance with relevant institutional and national guidelines and regulations. Animal housing and husbandry were maintained under the institutional holding permit granted by the local veterinary authority of the City of Bonn (Stadt Bonn) under reference number [67-5/06/2025] and the City of Essen (Stadt Essen) under reference number [32-2-1180-71/328]. All animal experiments were approved by the Landesamt für Verbraucherschutz und Ernährung (LAVE, permits AZ: 81-02.04.2017.A075 and AZ: 81-02.04.2023.A166) NRW, Germany, and conducted in accordance with the European Union Directive 2010/63/EU on the protection of animals used for scientific purposes.

### Brain template generation

To create an unbiased population template of the mole-rat brain, an established computational pipeline (*53–55*) (Felder et al, 2025, zenodo; Sirmpilatze et al, 2025, zenodo; Sirmpilatze et al, 2025, biorxiv) was used. The pipeline standardised the image orientation, corrected for intensity inhomogeneities, and automatically created masks of the brains. It then fed the images and masks to an existing, optimised implementation (*56*) of the symmetric group-wise normalisation (SyGN) algorithm (*57*) to create a brain template. SyGN iteratively registers and averages input images to the current best estimation of an unbiased template (Supplementary Figure 2).

We made two minor modifications to our pipeline to suit the mole-rat data. Firstly, the parallelisation tool “qbatch” (*58*) was integrated into the final template creation step, enabling distribution of sample registrations across 12 compute nodes on our institutional high-performance computer. This significantly sped up the pipeline. Secondly, to avoid excessive deformation of the mid-sagittal sections during non-linear registration, images of hemispheres were mirrored along their mid-sagittal plane during preprocessing, creating symmetric whole-brain images to pass to the SyGN algorithm.

### Magnetic field stimulation setup

Two custom-built triple-wrapped magnetic coil systems (Claricent, Germany) were used to generate controlled magnetic fields for the c-Fos and behavioural experiments. In the c-Fos experiments, we used a three-axis Helmholtz coil system placed inside a grounded Faraday cage. The magnetic coil dimensions were 61 cm (z-axis), 63 cm (x-axis), and 69 cm (y-axis). The coils were electrically shielded to minimise electric-field emission and powered by two low-ripple programmable power supplies (HMP4040 and HMP4030, Rohde & Schwarz Hameg, Germany). One of the three coil wraps represented the compensatory coil used to cancel the ambient magnetic field, while the other two wraps, the stimulation coils, either produced (with parallel current flow, CMF condition) or did not produce (anti-parallel, HMF condition) a changing magnetic field. This ensured that the current flow, and thus also possible side effects such as heat production and vibrations, were identical in the two magnetic conditions. The magnetic fields in this setup were measured with a one-axis fluxgate magnetometer (Fluxmaster, Stefan Mayer Instruments, Germany; range 1 nT to 200 μT, with a resolution of 1 nT). The residual field under HMF was reduced to below 1 μT.

The place preference experiments were performed inside a 3 m × 3 m × 2.5 m HF module cabin made of aluminium (Schwarz Shielding Systems, Germany). The cabin consists of 3 mm thick aluminium modules acting as a Faraday cage and additional insulation against sound. In the centre of the room, a heavy aluminium floor plate decoupled from the rest of the floor minimised vibration transmission to a non-magnetic table on which the animals were tested. All electronic lines entering the room were passed through low-pass filters to remove high-frequency components. Inside the room, a triple-wrapped 3D Merritt coil system with outer dimensions of 200 cm × 200 cm × 200 cm (Claricent, Germany), powered by a voltage-output module (NI-9264, National Instruments, USA) feeding into custom-made low-noise bipolar power amplifiers (Claricent, Germany), which created a homogeneous (>99%) magnetic field in the behavioural arena. The single set of compensatory coils was used to cancel the ambient geomagnetic field and the stimulation coils were controlled in real time to modulate field strength according to the animal’s position within the arena. During sham conditions, the compensatory coils were switched off, and the stimulation coils were powered in an anti-parallel configuration; therefore, the animals were exposed to the local geomagnetic field, and the changes in driving current did not produce magnetic field changes in the virtual anomaly region (potential confounders such as heat, vibrations or ramping noise from the power supplies were identical in this condition). A lockable blinding box was used for the switches to change between parallel and anti-parallel modes. Magnetic field intensity was mapped using a high-accuracy 3D fluxgate recording magnetometer (FGM3D TD, Sensys, Germany) equipped with an FGM3D/250 sensor. Furthermore, before each experiment, the coils were calibrated with a 3D fluxgate magnetometer (FVM400, Meda, USA). The coils were controlled and calibrated before each experimental session using custom Python scripts.

### Magnetic field exposure protocol (c-Fos experiment)

For the c-Fos experiments, mole-rats were placed individually in a round polycarbonate chamber (diameter = 20 cm; height = 19.8 cm) at the centre of the spherical Helmholtz coil system in the dark (IR-illumination, Fig. 3), covered with a perforated plexiglass sheet. To habituate the animals to the environment, they were placed in this arena in Earth’s magnetic field background for 30 min daily over the course of three days before the experiment. On the experimental day, the animals were placed in the chamber for 82 min. For the first ten minutes, all animals were exposed to a baseline HMF condition before the stimulation was started. In the CMF condition, the stimulation coils were configured in parallel and produced a dynamically varying field ranging between 20 and 70 μT with randomised direction changes every 30 s. The field stimulation consisted of two modes that alternated every 5 min: Mode 1: 90-degree directional changes every 30 s; Mode 2: Random magnetic field directions and intensity every 30 s (stimulus as in (*5*)). In the HMF condition, the stimulation coils were configured anti-parallel after compensation of the local geomagnetic field by the compensatory coil, yielding a near-zero (< 1 µT) residual magnetic field.

Animals were randomly assigned to CMF or HMF groups; two independent experimental replicates were performed in different years by different experimenters (Experiment A: CMF *n* = 5, HMF *n* = 6; Experiment B: CMF *n* = 5, HMF *n* = 4). Data from the two replicates were pooled for analysis to increase statistical power (total n = 10 animals per group). The animals from both experiments are labelled with open and filled circles, respectively, in all data plots.

### Brain tissue preparation and clearing

After the magnetic exposure, animals were deeply anaesthetised with ketamine and xylazine and transcardially perfused with heparinized phosphate-buffered saline (PBS) followed by 4% paraformaldehyde (PFA) in PBS. Brains were post-fixed in 4% PFA at 4 °C overnight and hemisected: The right hemispheres were processed for whole-brain clearing and immunostaining in this study, the left hemispheres were used in a separate study. Samples were processed according to the iDISCO+ protocol (*26*) with minor modifications. Briefly, tissues were dehydrated through a graded methanol series, bleached where indicated (skipped in tracer experiment), and permeabilised and blocked at 37 °C for 2 days per step (blocking/permeabilization buffer composition: (specify detergents, DMSO, glycine, serum)). c-Fos immunostaining was performed with rabbit anti–c-Fos antibody (Santa Cruz; catalogue no.: sc-253; dilution: 1:350) at 37 °C for 10 days with gentle rotation. Samples were washed and incubated with donkey anti-rabbit Alexa Fluor 647 secondary antibody (Thermo Fisher Scientific, catalogue no.: A-31573; dilution: 1:500) for 10 days. Following staining, samples were dehydrated, cleared in dichloromethane (DCM) and finally exchanged into dibenzyl ether (DBE) for refractive index matching and long-term storage. All incubations were performed in solvent-resistant polypropylene tubes with gentle rotation; care was taken to minimise oxidation during DBE storage by filling tubes completely and capping tightly.

### Light-sheet imaging

Cleared half-brains were imaged on a Miltenyi Ultramicroscope II (Miltenyi Biotec, Germany) light-sheet microscope. Imaging was performed using a (2x) objective (NA = 0.5) with a zoom setting of 0.8x, and excitation lasers at 488 nm (autofluorescence) and 647 nm (c-Fos). Image stacks were acquired with a z-step of 4 μm and a lateral sampling of 4.0625 μm per pixel; exposure times were set to 200 ms per plane and laser powers were adjusted per sample to avoid saturation. Image acquisition was controlled by Imspector Pro (version 7.4), and all data were exported as 16-bit TIFF stacks for further processing.

### Image processing and registration

Raw tiles were stitched using ClearMap 2.0 and Arivis (version 4.2) as needed. Each stitched brain volume was registered to an averaged mole-rat reference atlas at 20 × 20 × 20 μm isotropic resolution (available through the BrainGlobe Atlas API) (*29*) using Elastix (*59*) with an initial affine transform followed by non-linear B-spline registration (parameter settings as in (*26*)). Registration quality was assessed by visual inspection of anatomical landmarks across coronal, sagittal and horizontal planes; no registration was excluded from subsequent analyses.

### c-Fos quantification and p-value map generation

Detection of c-Fos–positive nuclei was performed in Ilastik (version 1.4.1b15) using a 3D pixel-classification model trained on manually annotated subsets covering representative regions of the brain (*60*). Probability maps were thresholded at *p* = 0.05 (probability cutoff) to generate binary detections; the resulting coordinates were exported as 3D point lists. Prior to density-map construction, raw detection coordinates were transformed into the standardised reference-brain coordinate space using the inverse registration transforms.

Voxel-wise density distributions were obtained by representing each detected nucleus as a 300-μm-diameter sphere of uniform intensity and summing these across the registered brain to yield an individual density map. Because the two experimental batches differed in total counts, voxel-wise statistical comparisons between CMF and HMF conditions were performed using a two-way ANOVA with condition and batch as factors. The raw voxel-wise *p*-value map (thresholded at p < 0.05) was subjected to cluster-based permutation correction: for each permutation iteration, registered cell coordinates were spatially randomised within the reference-brain mask and a two-way ANOVA was recomputed on the shuffled dataset. This permutation procedure was repeated 2,000 times to generate a null distribution of cluster sizes, and critical cluster-size thresholds were determined separately for clusters with increased density in CMF or HMF.

### Back-projection of significant clusters and post hoc validation

Significant clusters identified after cluster-correction were back-projected into each animal’s raw-image coordinate system using the inverse deformation fields computed during Elastix registration (rigid/affine/B-spline sequence as above). Back-projected cluster masks were dilated by 10 iterations of binary dilation (corresponding to ∼200 μm radial expansion) to compensate for minor registration variability and to ensure inclusion of neighbouring labelled nuclei. The dilated masks were overlaid on each animal’s raw c-Fos detections, and the number of nuclei within each dilated region was counted. These per-animal counts were used for post hoc comparisons between CMF and HMF groups to validate that atlas-level clusters corresponded to genuine differences in raw c-Fos signal.

### Behavioural analysis: c-Fos assay

Based on video recordings of the animals during the magnetic stimulations, a manual ethogram was created by a blinded observer (SWK) using the event logging software BORIS(*61*). We distinguished between four broad behavioural categories: walking, escaping/digging, resting, and other behaviours. We only annotated the first 40 min of the videos, as, due to the delay in expression, this reflects the timeframe in which most of the c-Fos that we measured was induced (*62*). We divided the total trial time (40 min: 10 min baseline + 30 min stimulation) into 5-minute bins and calculated the relative time spent in each behaviour category during that behaviour for each animal. Next, we calculated the relative time spent in each of the 4 categories during the baseline and stimulation periods, and subtracted the baseline from the stimulation for each animal.

### Closed-loop behavioural paradigm (place preference test)

Mole-rats were released into a linear track (length 80 cm) with semicircular ends placed at the centre of the Merritt-coils. Each end contained a non-magnetic novel object. A closed loop between the animal tracking and coil output resulted in a continuous adjustment of the magnetic environment in the arena, such that field intensity depended on the instantaneous animal position. The arena orientation within the room (major axes) was balanced across trials. Each trial lasted 20 minutes. A total of n = 15/16 animals were used per experiment, and no animal was tested twice. The study included three distinct behavioural experiments (EMF vs HMF; EMF vs Sham (anti-parallel but constant total current); EMF vs 2×EMF). Sex and trial order were counterbalanced across days to minimise order effects. The room was lit with 880 nm IR LEDs, and videos were recorded from above using an infrared camera (model: Basler camera, GigE acA1300-75gm; frame rate: 25 Hz; exposure time: 53.135 ms) for subsequent analysis.

Animal position was tracked using custom Python scripts by means of background subtraction, which detects the largest region of pixels that differ from the static background and records its centroid. Behavioural tracking data were downsampled to 5 Hz by analysing every 5^th^ frame of the 25 fps recordings. Spatial occupancy maps were computed as 2D histograms (10 × 50 or 50 × 10 bins depending on arena orientation) and smoothed with a Gaussian kernel (σ = 0.5 bins). Two lateral regions of interest (ROIs), each spanning 25% of arena width, were defined; histograms were rotated and aligned to ROI axes as needed. ROI occupancy fractions were computed as the fraction of frames within each ROI. An Avoidance Index was calculated as

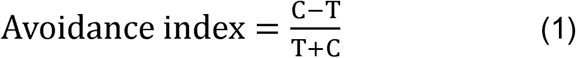

where *T* is the time spent in the test ROI (anomaly), and *C* is the time spent in the control ROI (resulting in values between 0 and 1, with 1 being absolute avoidance of the anomaly).

### Sensor measurement

To test the closed-loop magnetic field control in the place preference test, we moved a magnetic sensor through the linear track and measured the corresponding changes in field intensity. We used a SENSYS GmbH FGM3D/250 three-axis magnetometer, with a sampling rate of 400 Hz. We first placed the sensor at the arena centre, at the centre of the magnetic coils, then gradually moved it to one end designated as the virtual magnetic anomaly (HMF). As the sensor approached or moved away from the anomaly, the field intensity changed accordingly. Measurements were recorded continuously, and the sensor was held for approximately 10 s at both the anomaly and the centre. This procedure was repeated 10 times to obtain 10 measurements. We extracted the mean, standard deviation, and range from all sensor measurements obtained within a 5 cm diameter region at the arena centre and at the virtual magnetic anomaly.

### Statistics

We performed voxel-wise statistical comparisons to generate the whole-brain *p*-value map in customised ClearMap using a voxel-wise two-way ANOVA with magnetic condition (CMF vs HMF) and experimental batch (Experiment A vs B) as factors. To isolate magnetic-condition–related effects while controlling for batch variability, only the main effect of magnetic condition was used to compute voxel-wise *p*-values. This approach removes batch-driven differences by explicitly modelling batch as a second factor, thereby enabling the contrast of interest (CMF vs. HMF) to be tested independently of batch effects. The resulting raw voxel-wise *p*-value map was thresholded at *p* < 0.05 and subjected to cluster-based permutation correction (2,000 permutations) as described above.

To compare ethograms across magnetic field conditions, we used a general linear model with condition (CMF or HMF), behaviour, and their interaction as independent variables, and behavioural difference as the dependent variable. A post-hoc Tukey’s test was used to test for behavioural differences between the groups across the behaviours. To test whether animals changed their behaviour from the baseline to the stimulation period, we first performed a Shapiro-Wilk normality test for each group, with a Holm-Bonferroni correction for multiple comparisons. As *p* > 0.05 for each group, we performed a Student’s t-test to compare the mean of each group to 0, again with Holm-Bonferroni correction for multiple comparisons. Lastly, we performed a Principal component analysis (PCA), using the PCA function from the scikit-learn package(*63*).

All remaining statistical analyses were conducted in GraphPad Prism (v10.6.1) and R (v4.4.3). Total c-Fos–positive cell counts between CMF and HMF groups were compared using the Mann–Whitney test (two-tailed). Region-level comparisons of c-Fos densities were performed using the Mann–Whitney test, with multiple comparisons corrected using the False Discovery Rate (FDR) method (Q = 5%). Statistical significance is indicated as: *p_adj_* < 0.05 (*), *p_adj_* < 0.01 (**) and exact *p_adj_* values are reported in the main text. For behavioural tests, normality of the paired differences and the avoidance indices was assessed using Shapiro-Wilk tests and QQ plots. Occupancy differences between control and test regions were assessed using paired two-tailed *t*-tests. The avoidance index was evaluated using one-sample *t*-tests, each comparing the avoidance index against 0 (two-tailed). Statistical significance was defined as *p* < 0.05 unless otherwise specified. Exact n values and tests used are provided in the corresponding figure legends.

## Data and code availability

The raw and processed imaging datasets supporting the findings of this study are available from the corresponding author upon reasonable request. Custom code for image processing, registration, Ilastik/ClearMap integration, permutation testing, and behavioural analysis is available from the corresponding author upon reasonable request and will be made publicly available in a GitHub repository after publication. The mole-rat reference brain and corresponding annotations used in this study are publicly available (via the BrainGlobe Atlas API) (*29*). Additional materials are available from the corresponding author upon reasonable request.

## Acknowledgments

We acknowledge funding from the Max Planck Society, the European Research Council (ERC Starting grant no. 948728 to EPM), the Ministry of Culture and Science of the State of North Rhine-Westphalia (iBehave from the program “Netzwerke 2021”, grant no. NW21-049 to EPM) and the University Alliance Ruhr (MERCUR Starting Grant AN-2016-003 to EPM). We thank Georgina Fenton, Martha Daniel, and Christin Osadnik for discussions and help with data collection. Runita Shirdhankar received a doctoral scholarship from the DAAD and is supported by the International Max Planck Research School (IMPRS) for Brain and Behavior. Dori Grijseels gratefully acknowledges financial support from the Alexander von Humboldt Foundation through a Humboldt Research Fellowship. We are indebted to the animal unit and core facilities at the MPINB, and the IMP Vienna and David Keays for access to the light-sheet microscope.

## Author contributions

Conceptualisation: EPM

Methodology: LZ, AAF, NS, ALT, EPM

Validation: LZ, EPM

Formal Analysis: LZ, BYKL, AAF, NS, ALT, AS, LW, DG, EPM

Investigation: LZ, BYKL, AS, LW, AAF, NS, ALT, BH, RNS, SWK, DG, NA, SB, EPM

Data Curation: LZ, AAF, NS, ALT, EPM

Writing (Original Draft): LZ, EPM

Writing (Review and Editing): LZ, BYKL, AS, LW, AAF, NS, ALT, BH, RNS, SWK, DG, NA, SB, EPM

Visualisation: LZ, BYKL, AAF, NS, ALT, RNS, DG, EPM

Resources: SB, EPM

Supervision: EPM

Project Administration: EPM

Funding Acquisition: EPM

## Supplementary materials

**Supplementary Figure 1.**
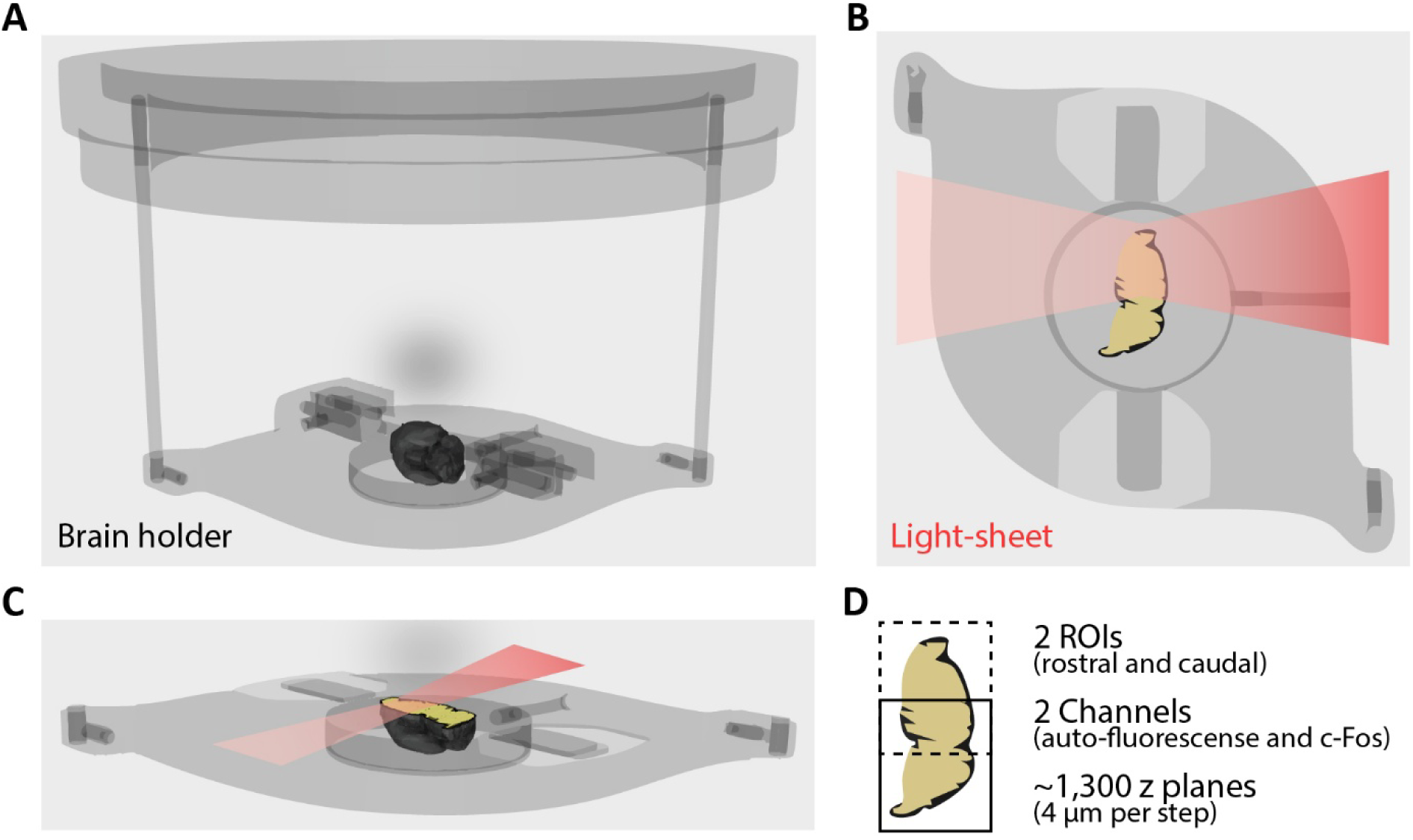
| Brain holder and light-sheet imaging. (**A**) A customised mole-rat brain holder with the processed brain positioned at the centre for light-sheet imaging. (**B**) A laser light sheet (red) illuminates the sample to generate optical sections in the sagittal plane. (**C**) A three-dimensional side view of the holder under light-sheet illumination. (**D**) Positioning of the two imaging FOVs that cover the entire brain.

**Supplementary Figure 2.**
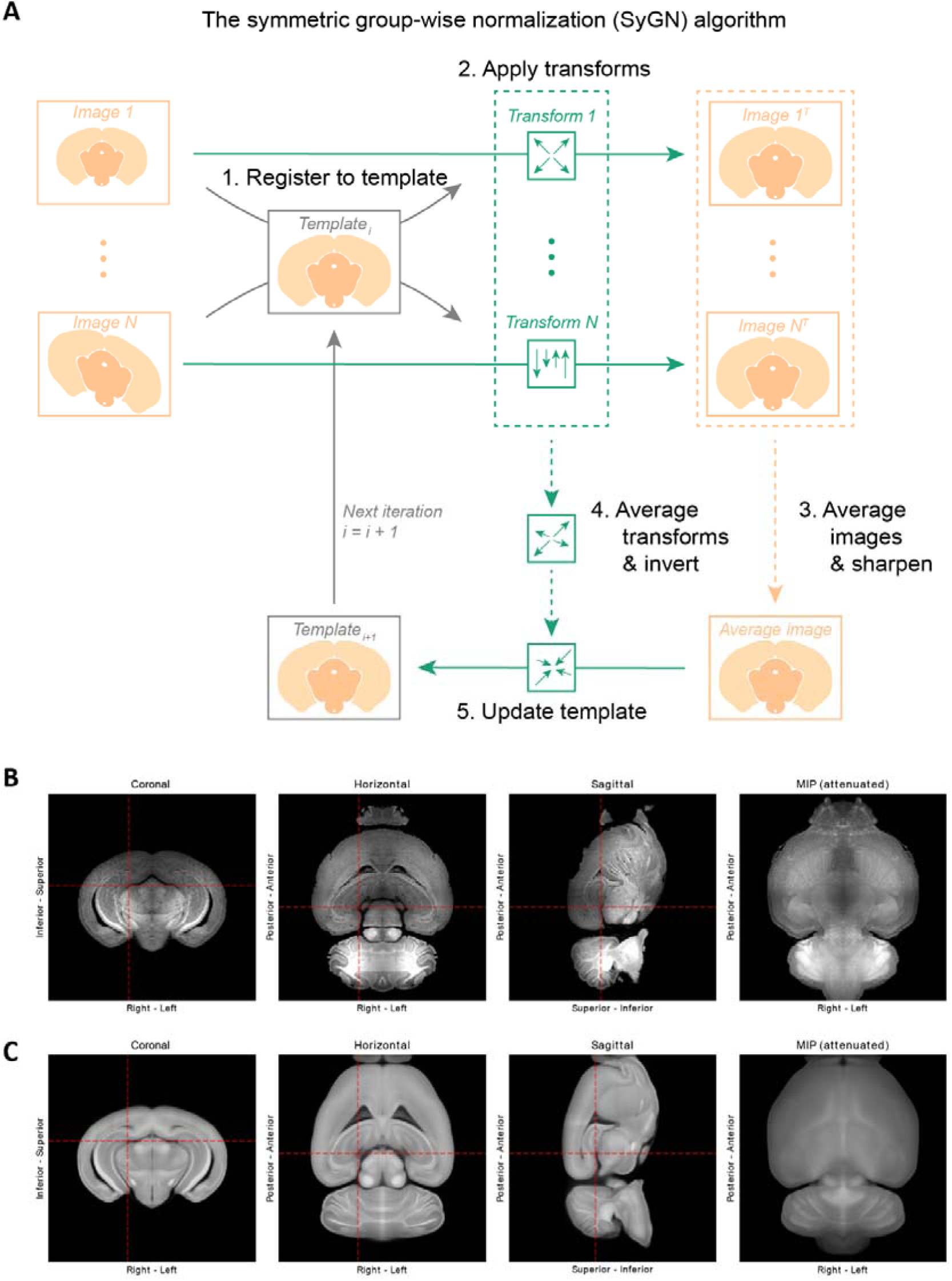
| Strategy of creating a mole-rat reference brain. **(A)** A schematic of the symmetric group-wise normalisation (SyGN) algorithm. Input images were created by mirroring hemispheric mole-rat brain acquisitions. SyGN uses rigid, similarity, affine and non-linear image registrations to iteratively converge on an unbiased average of its inputs. **(B)** Matching orthogonal sections of an individual brain compared to **(C)** our reference brain. This shows how sample damage and unique traits of the individual (e.g. the location of capillary blood vessels) are averaged out in the template, while anatomical features shared by all samples are preserved.

**Supplementary Figure 3.**
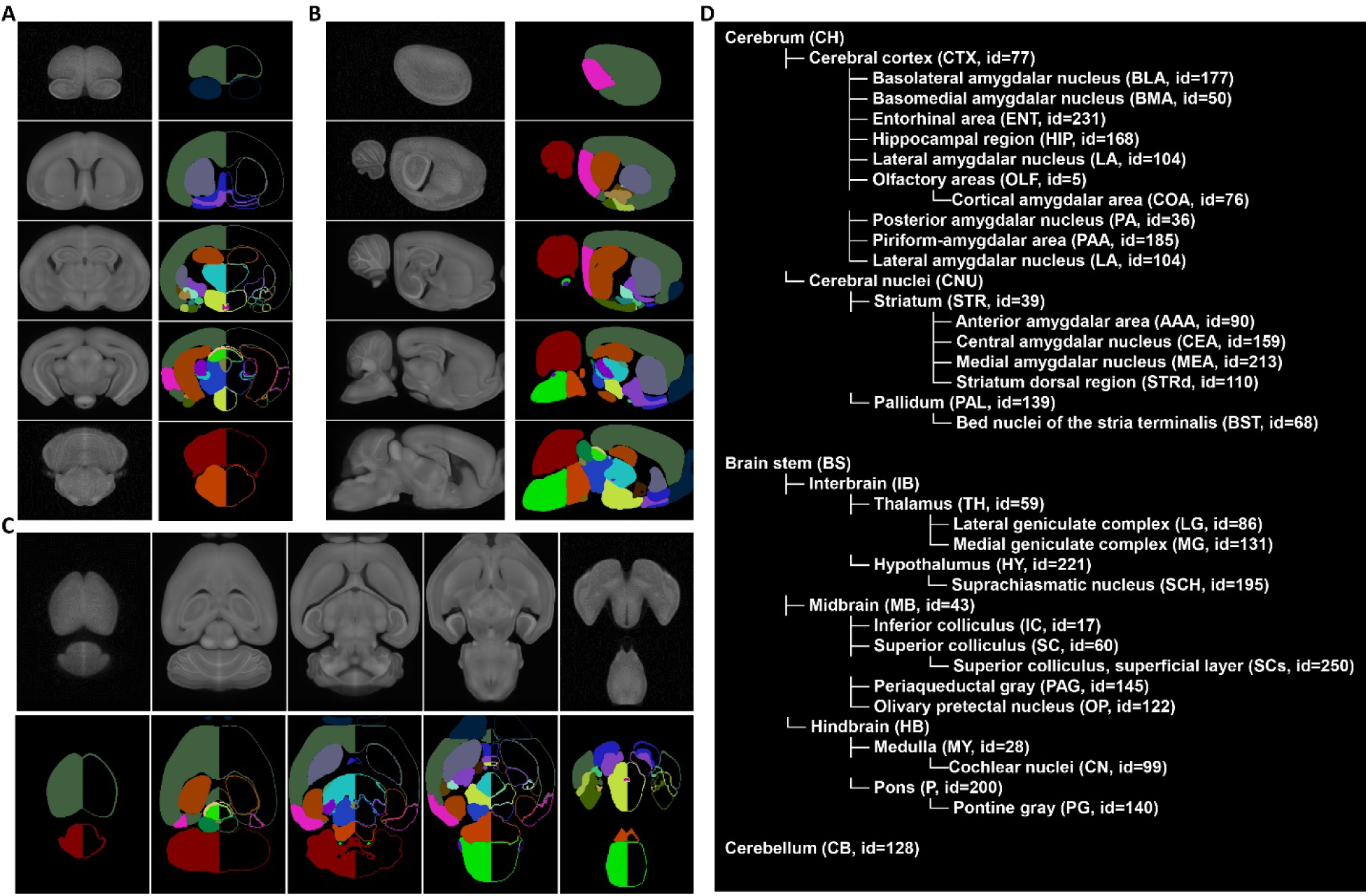
| Overview of the mole-rat reference brain annotations. (**A–C**) Coronal (A), sagittal (B), and horizontal (C) series showing the reference brain with corresponding annotations. Annotation boundaries are also indicated in (A) and (C). (**D**) Hierarchical tree representation of annotated brain structures.

**Supplementary Figure 4.**
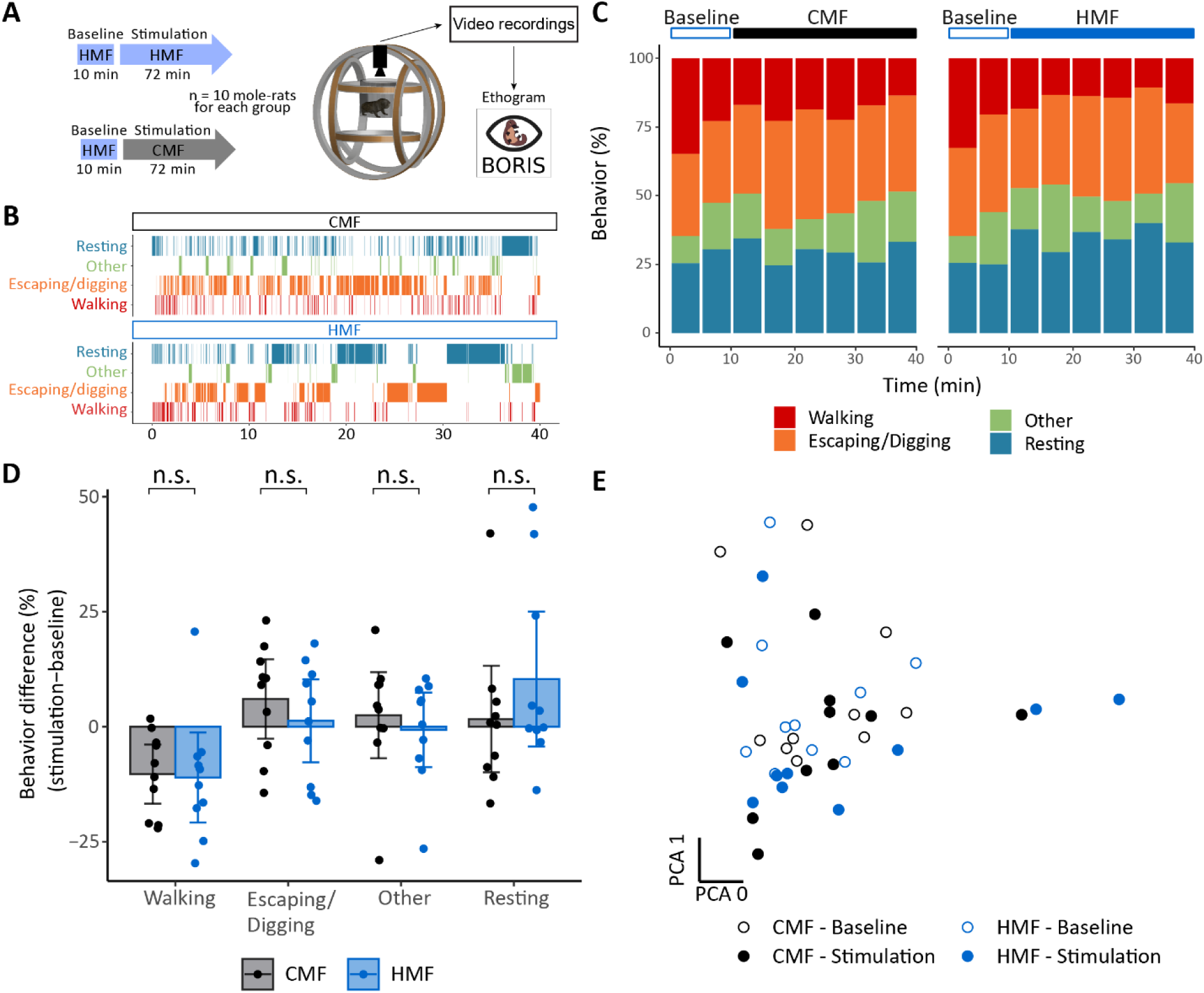
| No behavioural differences in mole-rats during magnetic field exposure. (**A**) Experimental paradigm. Video recordings were used to create an ethogram in BORIS software by an observer blinded to the treatment conditions. **(B)** Example ethograms of two animals. **(C)** Average fractions of observed behaviours of all animals in both treatment groups over time (5-minute bins). **(D)** Behaviours during the stimulation phase normalised to the baseline phase indicated that walking decreased over time, but there was no difference between the two treatment groups for all scored behavioural categories. **(E)** Principal component analysis showed no clustering of the treatment groups for the first two principal components, which together explained 86.3%% of the variance.

**Supplementary Figure 5.**
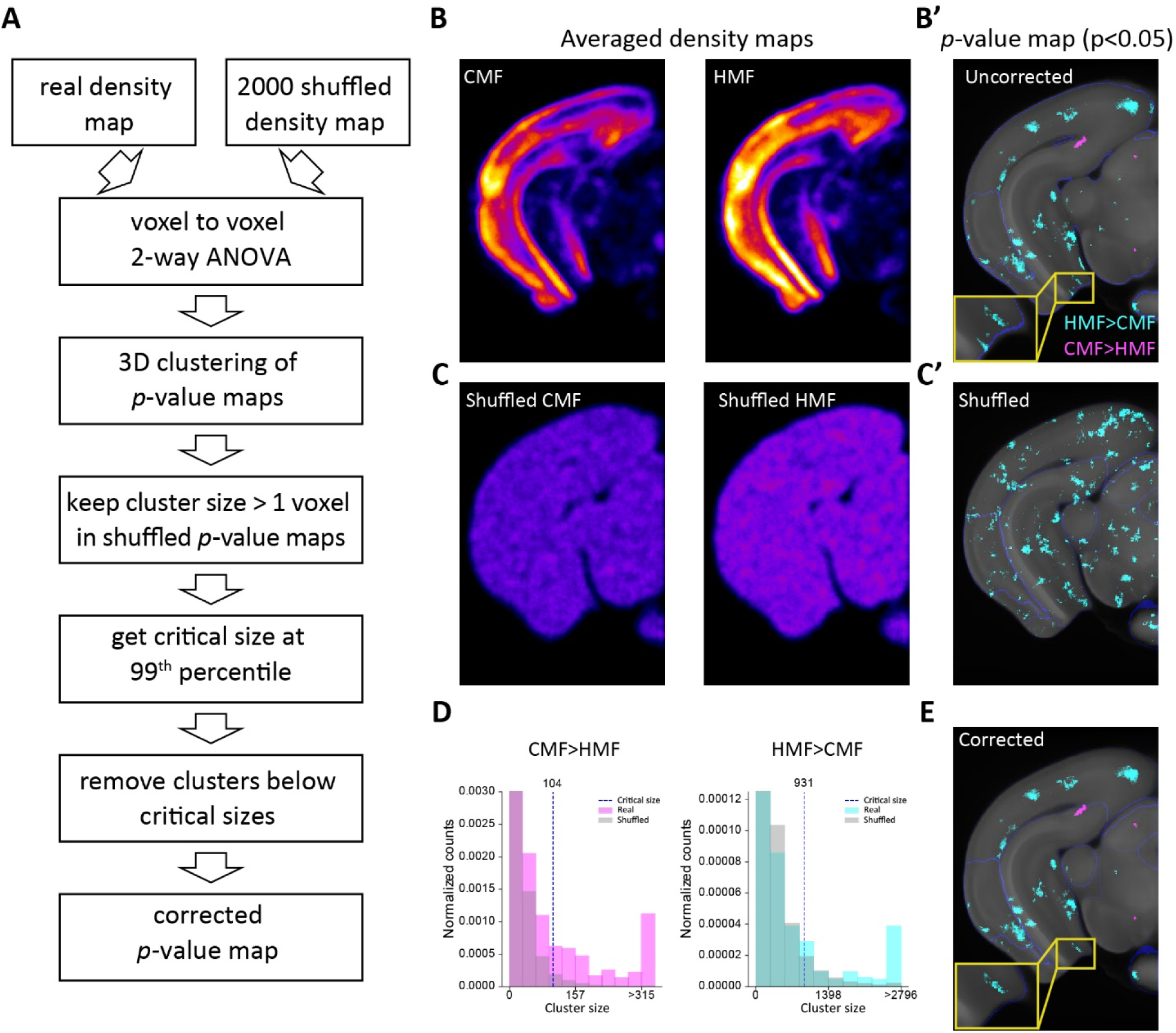
| Permutation-based correction of the *p*-value map for multiple testing. (**A**) Schematic of the cluster-based correction procedure. (**B**) Representative planes of real data showing cell density maps (left and right) and the uncorrected *p*-value map (B′). (**C**) Corresponding visualisation for a shuffled simulation example and the simulated p-value map (C′). (**D**) Histogram of cluster size distributions for decreased (left) and increased (right) activation in the HMF group (2,000 shuffled iterations); the dotted line indicates the critical cluster-size thresholds. (**E**) The same plane as in (B’) and (C’) for the corrected *p*-value map after applying the thresholds. Yellow boxed insets in (B’) and (E) show clusters that were below the threshold and removed after correction.

**Supplementary Figure 6.**
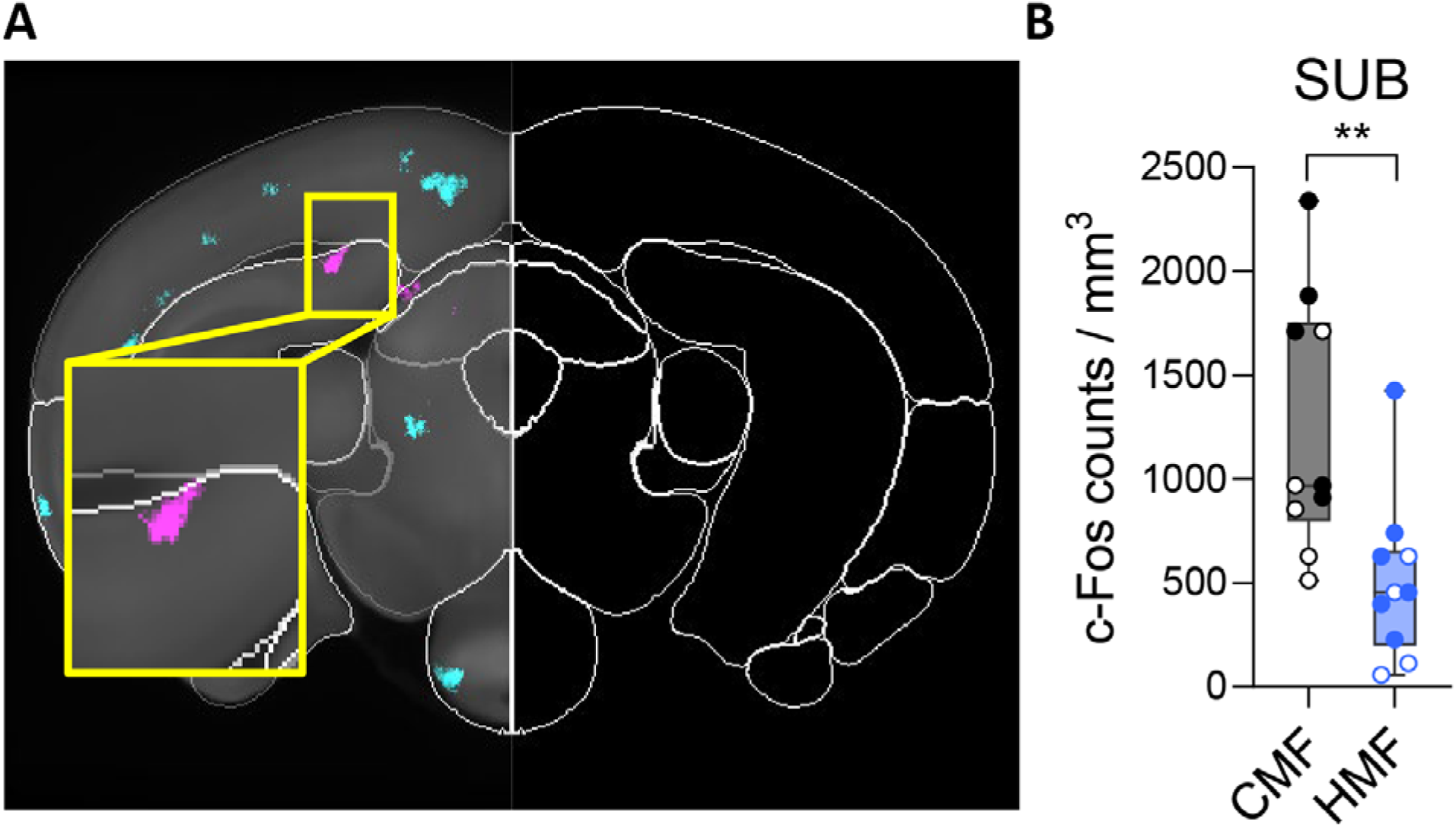
| Changing magnetic fields increase neuronal activity in the dorsal subiculum. (**A**) A single plane of voxel-wise p-value maps with an inset. (**B**) Box plots comparing cell densities; individual data points are shown. n = 10 mole-rats per group; solid and open data points represent two independent experiments. Statistical significance was assessed using a Mann–Whitney test with FDR correction (Q = 5%), and exact *p_adj_* values are reported in the main text. ** indicates *p_adj_*< 0.01. SUB: subiculum.

**Supplementary Figure 7.**
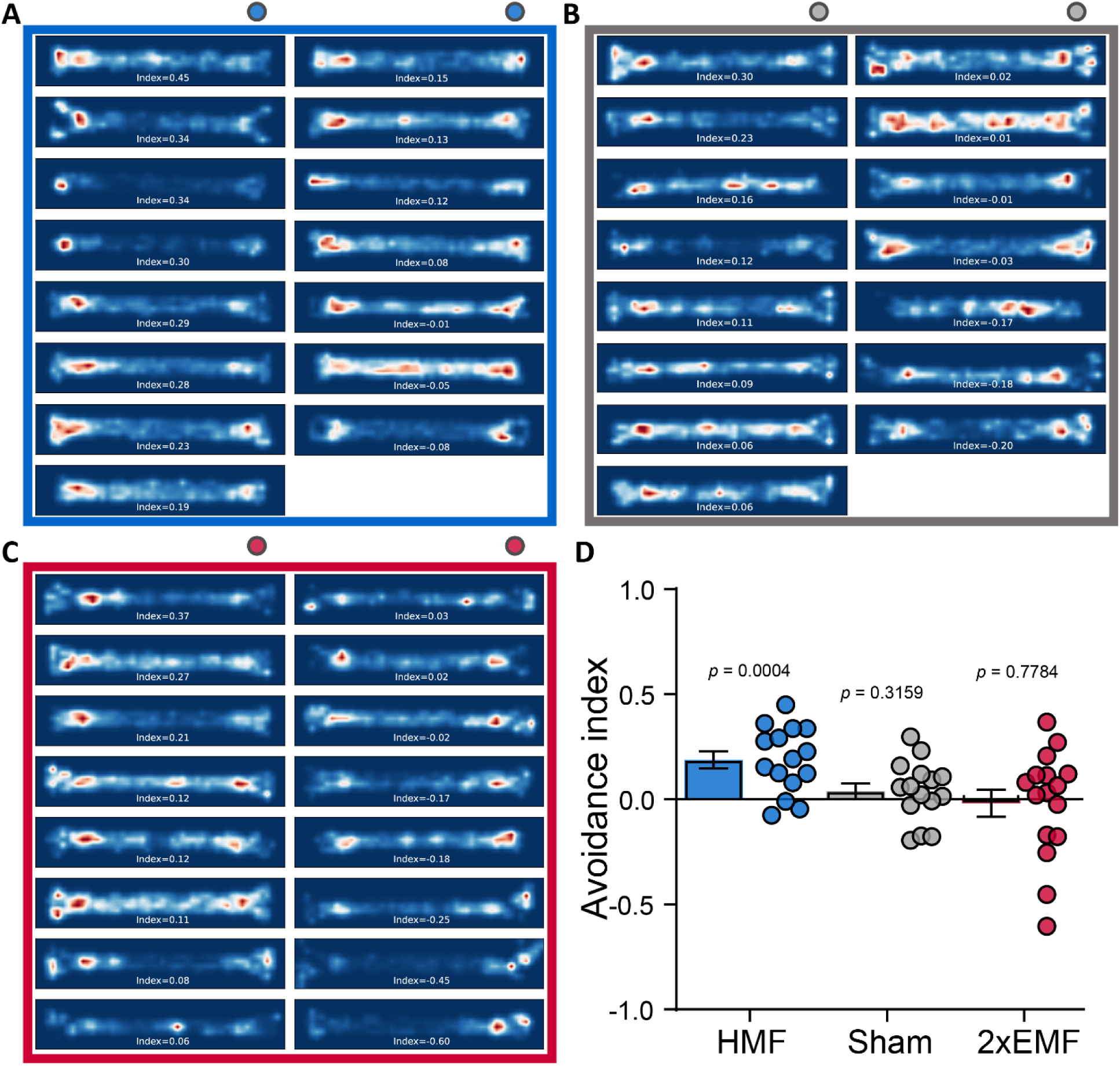
| Individual occupancy heatmaps and avoidance index from the closed-loop virtual magnetic field tests. (**A–C**) All individual occupancy heatmaps across three experimental conditions: HMF (A), Sham (B), and 2×EMF (C), in order of descending avoidance indices. The dots indicate the end of the virtual anomaly (always shown on the right). (**D**) Avoidance indices in the three experimental conditions. Statistical difference from 0 was assessed by one-sample t-tests: EMF vs HMF, *t* (14) = 4.635, *p* = 0.0004; EMF vs Sham, *t* (14) = 1.040, *p* = 0.3159; EMF vs 2×EMF, *t* (15) = –0.287, *p* = 0.7784.

**Supplementary Table 1.**
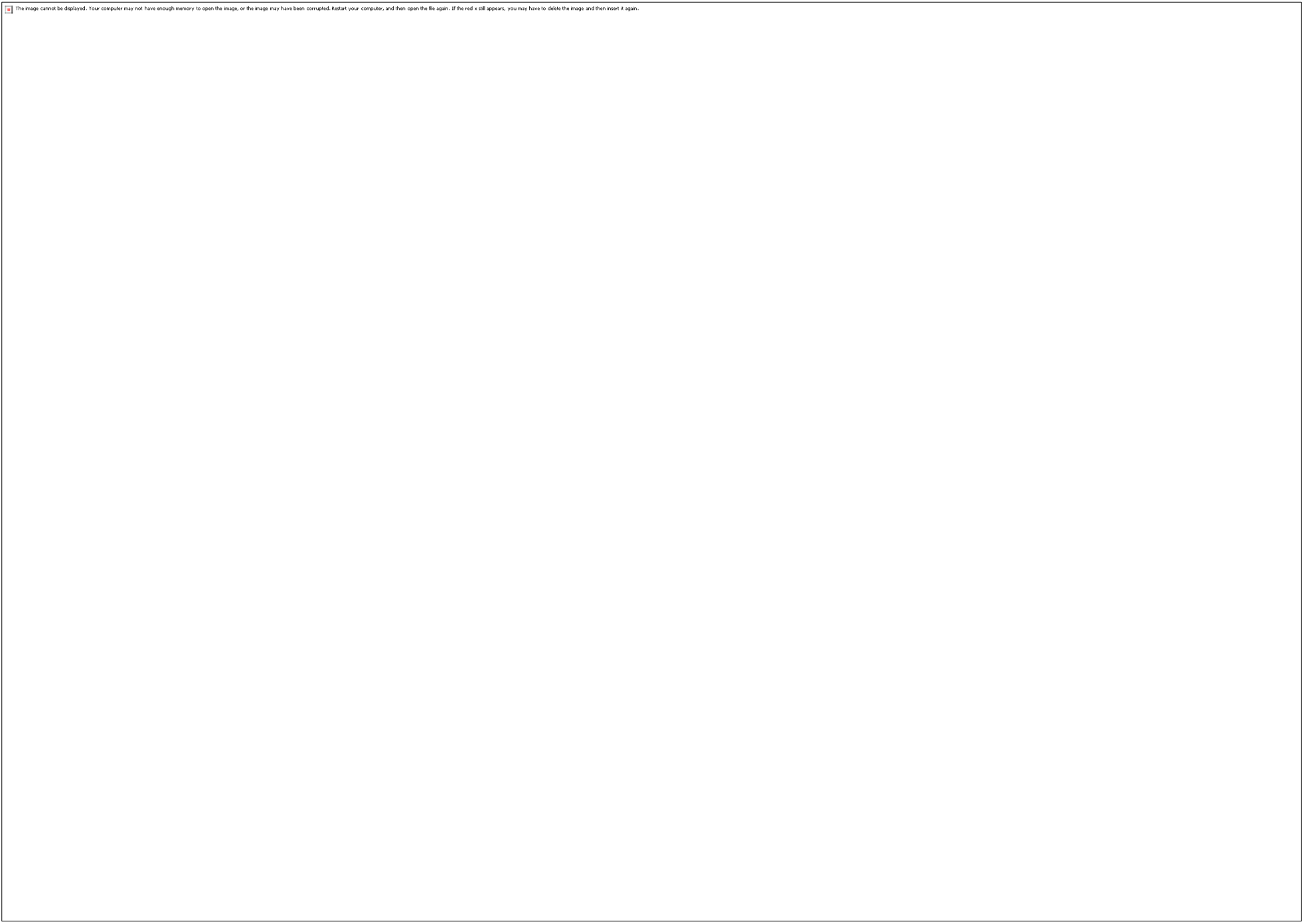
| *p*-value map significant voxel counts based on annotation atlas.

**Supplementary Table 2.**
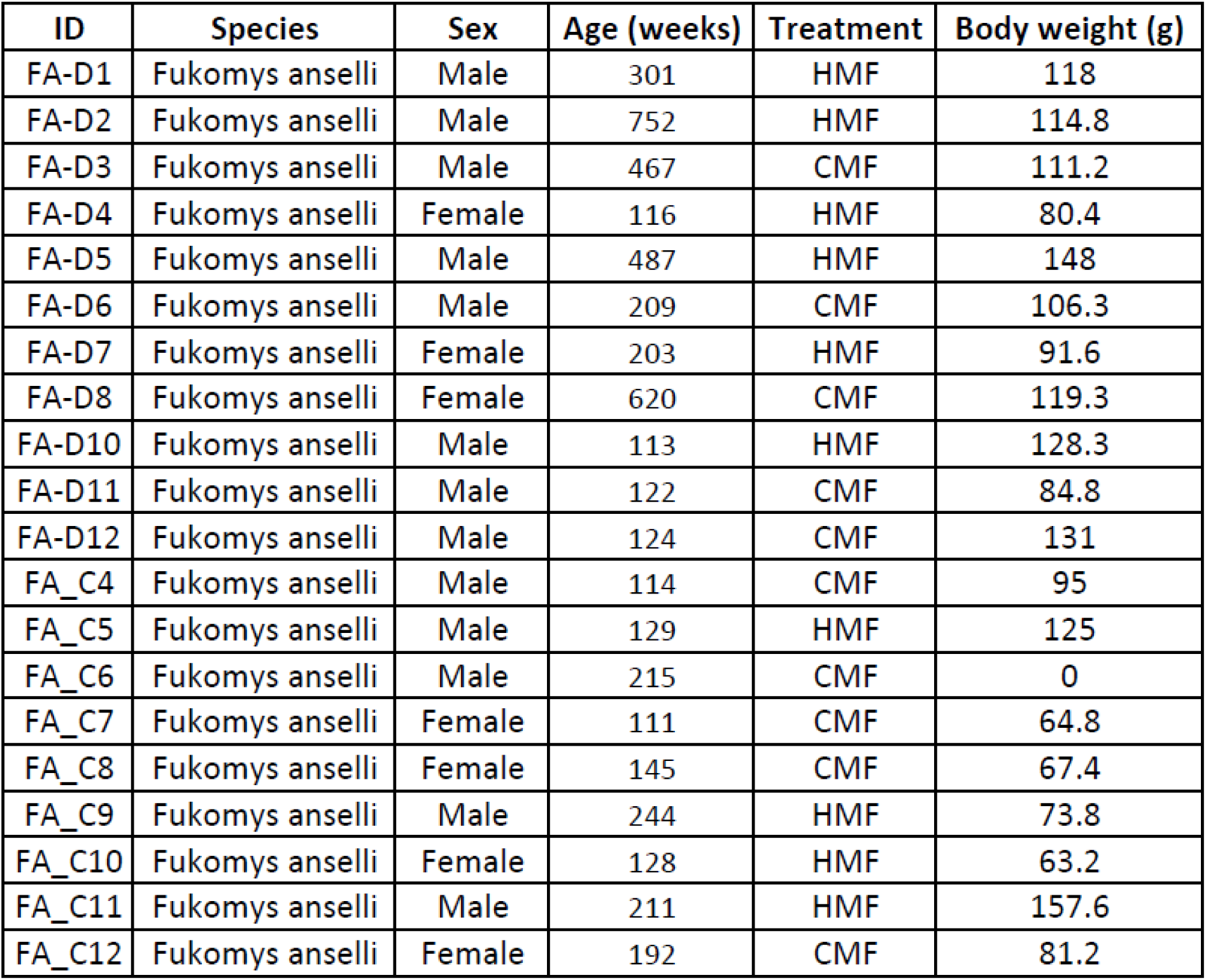
| animals used in c-Fos experiments.

